# Defining minimal selective concentrations of amoxicillin, doxycycline and enrofloxacin in broiler-derived cecal fermentations by phenotype, microbiome and resistome

**DOI:** 10.1101/2023.11.21.568155

**Authors:** Aram F. Swinkels, Egil A.J. Fischer, Lisa Korving, Nina E. Kusters, Jaap A. Wagenaar, Aldert L. Zomer

## Abstract

Antimicrobial resistance (AMR) is an emerging worldwide problem. Exposure to antimicrobials selects for resistant bacteria which are a health threat for humans and animals. The concentration at which selection for resistant bacteria occurs is often lower than the minimum inhibitory concentration (MIC) and also differs between environments. Defining this minimal selective concentration (MSC) under natural conditions is essential to understand the selective window for resistant bacteria which are exposed to residual antimicrobials in humans, animals and the environment.

In this study we estimated the MSCs of three antimicrobial compounds, amoxicillin, doxycycline and enrofloxacin in a complex microbial community by conducting fermentation assays with cecal material derived from broilers. We examined the phenotypic resistance of *Escherichia coli*, resistome and microbiome after 6 and 30 hours of fermenting in the presence of antimicrobials of interest. The concentrations are 10 to 100 times lower than the epidemiological cut-off values in *E. coli* for the respective antimicrobials as determined by EUCAST (https://mic.eucast.org/). In contrast to the amoxicillin and doxycycline exposure we could not determine any molecular resistance mechanism in the resistome analysis for the enrofloxacin exposure, likely because they are the result of point mutations.

Our findings show at which concentrations there still is selection for AMR bacteria. This knowledge can be used to manage the risk of the emergence of AMR bacteria.

**Importance:** Antimicrobial resistance is an emerging threat to the health of humans and animals; it might affect economic prosperity in the future. The rise of antimicrobial resistant bacteria is a consequence of the use of antimicrobial compounds in humans and animals which selects for resistant bacteria. It is critical to understand the relation between the concentrations of antimicrobial compounds and their selection for antimicrobial resistant bacteria. In our study we are providing the minimal selective concentrations for amoxicillin, doxycycline and enrofloxacin by using cecal fermentations assays.

## Introduction

It has been estimated that in 2050 more deaths will be caused by resistant bacteria than cancer if no measures are taken (1). This is mainly driven by the excessive use of antimicrobials in human medicine and in agriculture (2). This (mis)use of antimicrobial compounds has led to an alarming situation where resistant bacteria are a threat to the health of humans and animals (3). In addition, discovery of new antimicrobial compounds is going slow which is problematic since their powerful properties as medicines (4, 5). Despite all successful efforts to decrease antimicrobial usage in the last couple of decades, it is imperative to continue research on how and when selection of AMR bacteria takes place. An understudied field in relation to AMR are antimicrobial residues which remain in the environment long after their application time.

In livestock, normally, antimicrobial treatment lasts several days followed by a withdrawal period in which it is assumed that the antimicrobial compound including its activity disappears (6, 7). However, some antimicrobials are able to remain longer in the animal or environment due to stable half-life properties (8, potentially resulting in an extended period of selection for AMR bacteria. The minimal inhibitory concentration (MIC) is the lowest concentration of an antimicrobial compound that can inhibit the growth of bacteria. In pharmacodynamic models it is generally assumed that selection of AMR bacteria with an antimicrobial only occurs between the MIC of a susceptible bacterium and the MIC of a resistant bacterium, mainly due to the fitness costs of harboring a resistance gene or mutations (9, 10). On the contrary Gulberg et al (2011 described that sub-MICs, concentrations ranging between the minimal selective concentration (MSC) and the MIC of susceptible bacteria, can also select for AMR bacteria(11, 12). The minimal selective concentration is the lowest concentration which favors the growth of AMR bacteria over antimicrobial susceptible bacteria (12).

Selection for AMR bacteria occurs mostly in the gastrointestinal tract as conditions are favorable for outgrowth of resistant bacteria due to conditions that promote bacterial growth and niche competition (13, 14). Besides this, the high density of bacteria facilitates horizontal gene transfer between organisms in the gut microbiome as well as the increased chance of *de novo* resistance mutations because of the high cell densities (1, 15). Therefore, the concentration of antimicrobials reaching the gut plays a crucial role in the selection for AMR bacteria as resistant bacteria can out-compete the susceptible bacteria influenced by the selective pressure of the antimicrobial present at residual concentrations (16). For example, farm animals can excrete the antimicrobial compounds partially or in an unchanged biochemical structure which might result in re-exposure to antimicrobials in the gastrointestinal tract through coprophagic behavior (17). These residual concentrations may be high enough to exceed the MSC and therefore extend selection for resistant organisms long after treatment has ended. Another essential point is that some estimations of the MSC might be 10 to 100 times lower than the MIC of susceptible bacteria (11). The residual concentrations of a treatment can therefore affect the emergence of AMR and for that reason defining the MSC will provide important information.

The MSC has been determined in several other studies however they were mostly conducted in rich medium (18). This is in contrast to the gut environment where nutrients are more scarce as consequence of poor mixture and the flow through the gut (19). Besides, competition may be fierce between different bacterial species (20). Therefore, it is possible that antimicrobial compounds can select for AMR bacteria in the gut microbiome at different concentrations. By defining the MSC in non-model-organisms using e.g. resistome or microbiome measurements, it is possible to provide estimates for the selection of resistant bacteria in humans, animals or the environment.

The aim of this study was to determine the MSC of amoxicillin, doxycycline and enrofloxacin in a complex microbial community as found in poultry. Besides, we looked at the microbiome composition after exposure of the antimicrobial compounds. Therefore, we investigated this by conducting cecal fermentation assays by exposing cecal contents of broilers to sub-MIC levels of the selected antimicrobials to mimic the natural environment in the intestine to observe MSCs. At first, we determined the MSC by analyzing phenotypic resistance of *Escherichia coli* isolates in sub-inhibitory concentrations. This was done since *E. coli* is used as an indicator organism due to its simplicity to study in the lab and represents a significant fraction of the gut microbiome. In addition to selecting for phenotypic antimicrobial resistance of one species (*E. coli)*, sub-inhibitory concentrations can alter the microbial composition and changes after exposure are therefore an indication for the MSC. For that reason, we determined the microbial composition after exposure to the different antimicrobials to observe a change in the species diversity and presence. Besides the microbiome composition, an increase of resistance genes in the metagenome (the resistome) can reveal selective properties of the antimicrobial compound. Therefore, we also investigated the resistome for a possible increase in resistance genes after the treatment.

## Material and methods

### *In vitro* enriched competition assay

To estimate which concentrations select for AMR bacteria and use these in further experiments, we used isogenic *E. coli* strains in an enriched competition assay (Table 1). The stains were grown overnight in a liquid LB culture at 37 °C. After overnight incubation the susceptible strain and the resistant strain were mixed in a 1:4 ratio, this ratio was chosen in order to observe the possible increase of the resistant strains. From this mixture 3 μl was inoculated in a tube containing 3 mL LB medium and incubated for 6 hours at 37 °C. Besides a control sample, samples were treated with amoxicillin (0.08 mg/L, 0.8 mg/L, 8 mg/L and 80 mg/L), doxycycline (0.004 mg/L, 0.04 mg/L, 0.4 mg/L and 4 mg/L) and enrofloxacin (0.00125 mg/L, 0.0125 mg/L and 0.125 mg/L). Next, a 1:10 dilution in LB medium was prepared for each sample. From each 1:10 dilution five samples were taken using a 10 μl inoculating loop and inoculated on individual MacConkey agar plates. These plates were subsequently incubated overnight at 37 °C. Afterwards, the five plates per sample were used to select 96 individual colonies which were phenotypically confirmed to be *E. coli* which were each transferred to a different well of a 96 wells plate containing 100 μl of LB. The 96 colonies per treatment were transferred to squared MacConkey plates, containing epidemiology cut off values (ECOFFs) concentrations of amoxicillin (8 mg/L), doxycycline (4 mg/L), enrofloxacin (0.125 mg/L), cefotaxime (1 mg/L) and a control plate without antimicrobials using a stamp. An *E. coli* colony was scored non-wildtype (further called resistant) if it was able to grow on the plate with ECOFF concentration. This allowed us to calculate the percentage of resistance of the selected *E. coli’s* by comparing the growth of the *E. coli* on the control plate with the growth on the selection plates.

**Table 1.** Strains used in enriched competition assays.

| Strain | Source | Resistance | Used in enriched assay |
| --- | --- | --- | --- |
| **W3110** | Bachmann B. 1972 (21) | No resistance | Doxycycline and Amoxicillin |
| **W3110 + IncHI1 plasmid** | Transformation of W3110 in our lab with plasmid p12S05273-2 (22, 23) | Amoxicillin (MIC 16 mg/L)<br>Doxycycline (MIC > 16 mg/L) | Doxycycline and Amoxicillin |
| **37** | Isolated from a cecal fermentation experiment (1<sup>st</sup> repetition) | Amoxicillin (MIC > 8 mg/L)<br>Doxycycline (MIC > 4 mg/L) | Enrofloxacin |
| **37.3** | From strain 37 by performing a direct exposure assay (SNP difference 58) | Amoxicillin (MIC > 8 mg/L)<br>Doxycycline (MIC > 4 mg/L)<br>Enrofloxacin (MIC > 0.125 mg/L)<br>Flumequine (MIC > 2 mg/L) | Enrofloxacin |

### Sample collection and cecal fermentation

Caecum material was obtained from broilers from commercial farms in The Netherlands which were culled for welfare reasons as they encountered serious lameness. The broilers had an age varying from 15 to 40 days this was due to different sample moments as this experiment was repeated six times and for each experiment caecum material of two different broilers was used. The caeca were removed from euthanized broilers which were used for educational purposes at the faculty of veterinary medicine (registration-number: AVD10800202115056). The caeca were transported to the lab and within half an hour the contents were obtained from the caeca. Subsequently the cecal contents of the different caeca were mixed to get a homogenized mixture. Thereafter, 1% inoculation was prepared in 10 ml viande leuvere (VL) medium (24, 25) in a tube and 1 ml was added to an Eppendorf tube. The VL medium was modified by adding cysteine hydrochloride (0.4 g/L), 20 % glucose, vitamin K (9 mg/L), hemin (50 mg/L) and bile (5 mg/L) after adjusting the pH to 6.3 and autoclaving. In addition, to buffer the medium, sodium hydrogen phosphate (2.5 mg/L) was added before autoclaving. The fermentations were treated with amoxicillin (8 mg/L, 0.8 mg/L and 0.08 mg/L), doxycycline (4 mg/L, 0.4 mg/L and 0.04 mg/L), enrofloxacin (0.125 mg/L, 0.0125 mg/L and 0.00125 mg/L) and a control treatment was included. The incubation was done in a shaker at 41 °C at 300 rpm in an anaerobic hood to mimic the conditions in the intestine. At the beginning of the experiment a baseline measurement was conducted. Subsequently, a 6- and 30-hour measurement was taken for every treatment. This experiment was repeated 6 times and every experiment was performed on a different day as the caeca from different broilers from different flocks were used.

### *E. coli* isolation and phenotypic selection determination

Directly after removing the Eppendorf tubes from the anaerobic hood 10 µl of each fermentation was inoculated five times, on five individual MacConkey plates. After overnight incubation at 37 °C approximately 20 single colonies per MacConkey plate were picked with a pipette tip and transferred to a single well in a 96-wells plate that contained 100 µl LB medium per well. The 96 colonies per treatment were transferred, using a stamp, to squared MacConkey plates containing ECOFF (epidemiology cut off values) concentrations of amoxicillin (8 mg/L), doxycycline (4 mg/L), enrofloxacin (0.125 mg/L), cefotaxime (1 mg/L) and a control plate without antimicrobials. Next, resistant colonies were scored after overnight incubation at 37 °C. An *E. coli* colony was scored resistant if it was able to grow on the plate with ECOFF concentration of the specific antimicrobial. This allowed us to calculate the percentage of resistance of the selected *E. coli’s* by comparing the growth of the *E. coli* on the control plate with the growth on the selective plates.

### Shotgun metagenomics

DNA from the samples was extracted according to the EFFORT protocol and the DNA concentrations were measured with a Qubit (26, 27). Next, Illumina sequencing was performed using Illumina NovaSeq 6000 (Useq, Utrecht sequencing facility) with a maximum read length of 2 x 150 bp. Libraries were prepared with Illumina Nextera XT DNA Library Preparation Kit according to the manufacturers protocol (28). Afterwards, the reads were trimmed with trim galore (v0.6.4_dev) and the quality was assessed with FastQC (v0.11.4). Afterwards the reads were analyzed for the taxonomic classification by kraken2 and subsequently the abundance of the DNA sequences was computed with bracken (29, 30). The output of these packages was summarized into a biomfile using kraken2biom and were analysed with the R programme packages Phyloseq (v1.36.0), Microviz (v0.9.1) and Microbiome (v1.14.0) by which the species composition and alpha and beta diversity were estimated from a rarefied phyloseq object (31, 32). Finally, the resistome was investigated by using the tool KMA which detects alignments known to antimicrobial resistance genes using the Resfinder database (33, 34). The reads were first normalised for gene length and displayed as sequence depth per gigabase.

### Statistical analysis of the phenotypic resistance *E. coli* assay, resistome and microbiome of the caecal fermentations

We used a Poisson model for count data and a linear model for continuous data with the repetitions as random effect, using the glm function in R (4.1.0). We performed a post hoc Tukey test to determine whether the treatment groups at the time points 6 and 30 hours were significantly different from their control treatment. The R-script and data files can be found in (zenodo repository).

## Results

### Enriched media competition with isogenic *E. coli* strains

The resistant strain was able to out-compete the susceptible strain at a concentration of 8 mg/L for amoxicillin in rich media (Figure 1). For the doxycycline treatment a concentration of 0.4 mg/L the resistant strains had the advantage over the susceptible strain. The enriched competition with enrofloxacin showed a shift in resistant *E. coli* at a concentration of 0.0125 mg/L. The concentration found in the enriched competition give an indication of the minimal selective concentrations to be investigated in fermentation experiments.

**Figure 1.**
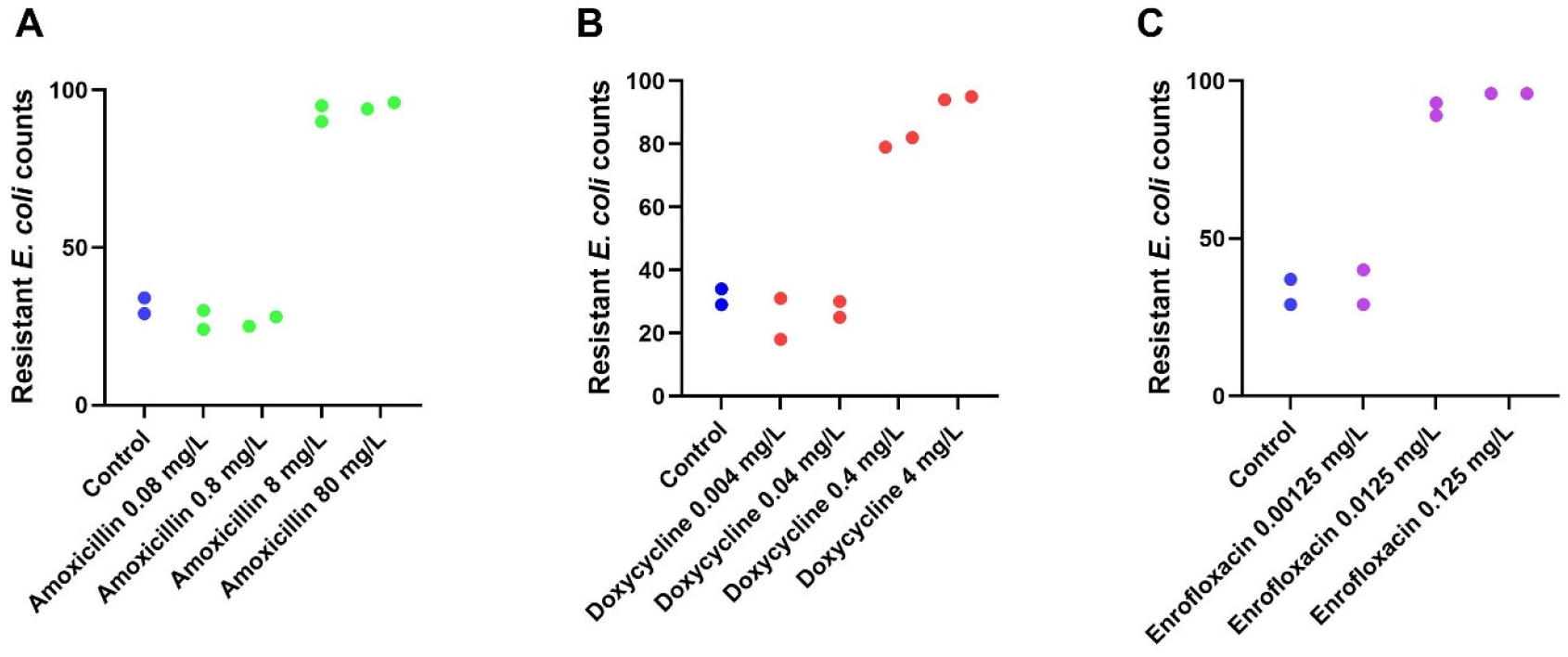
*Enriched competition assays. **A** shows the enriched competition with amoxicillin. **B** the enriched competition with doxycycline and **C** the competition with enrofloxacin.*

### Phenotypic resistance of *E. coli* isolates from caecal fermentations after exposure to antimicrobials

In the ceacal fermentations, we observed a clear shift in terms of resistant *E. coli* isolates for all the treatments which is shown in Figure 2. The amoxicillin treatment showed an increase in resistant *E. coli* isolates at 6 and 30 hours and treated with the concentration 0.8 mg/L or 8 mg/L compared to the control group (6 hours 0.8mg/L p<0.0001, 30 hours 0.8 mg/L p<0.0001, 6 hours 8 mg/L p<0.0001, 30 hours 8 mg/L p<0.0001). This indicates that the MSC of amoxicillin for *E. coli* ranges between 0.08 mg/L and 0.8 mg/L. The treatment with doxycycline showed that a concentration of 4 mg/L at timepoint 6 and 30 hours an increase in resistant *E. coli* compared to the control group (6 hours 4 mg/L p<0.0001, 30 hours 4 mg/L p<0.0001). Besides, the treatment with 0.4 mg/L doxycycline showed a significant value at 6 hours. However, we did not show this point in the graph since some experiments had limited to zero doxycycline resistant *E. coli* isolates. We performed the statistical tests without the experiments with a few or zero doxycycline resistant *E. coli* which resulted in a non-significant difference between the 6 hours 0.4 mg/L doxycycline sample and the control group at 6 hours in contrast to the treatments with 4 mg/L doxycycline. The MSC of doxycycline for *E. coli* is therefore estimated between 0.4 mg/L and 4 mg/L. Lastly, we observed an increase in the resistant *E. coli* isolates at a concentration of 0.125 mg/L enrofloxacin at timepoints 6 and 30 hours (6 hours 0.125 mg/L p<0.0001, 30 hours 0.0125 mg/L p=0.0005) also at a concentration of 0.0125 mg/L enrofloxacin at 30 hours (p<0.0001). Suggesting that the MSC of enrofloxacin for *E. coli* at 6 hours is ranging between 0.0125 mg/L and 0.125 mg/L. For the treatment at 30 hours it is estimated between 0.00125 mg/L and 0.0125 mg/L.

**Figure 2.**
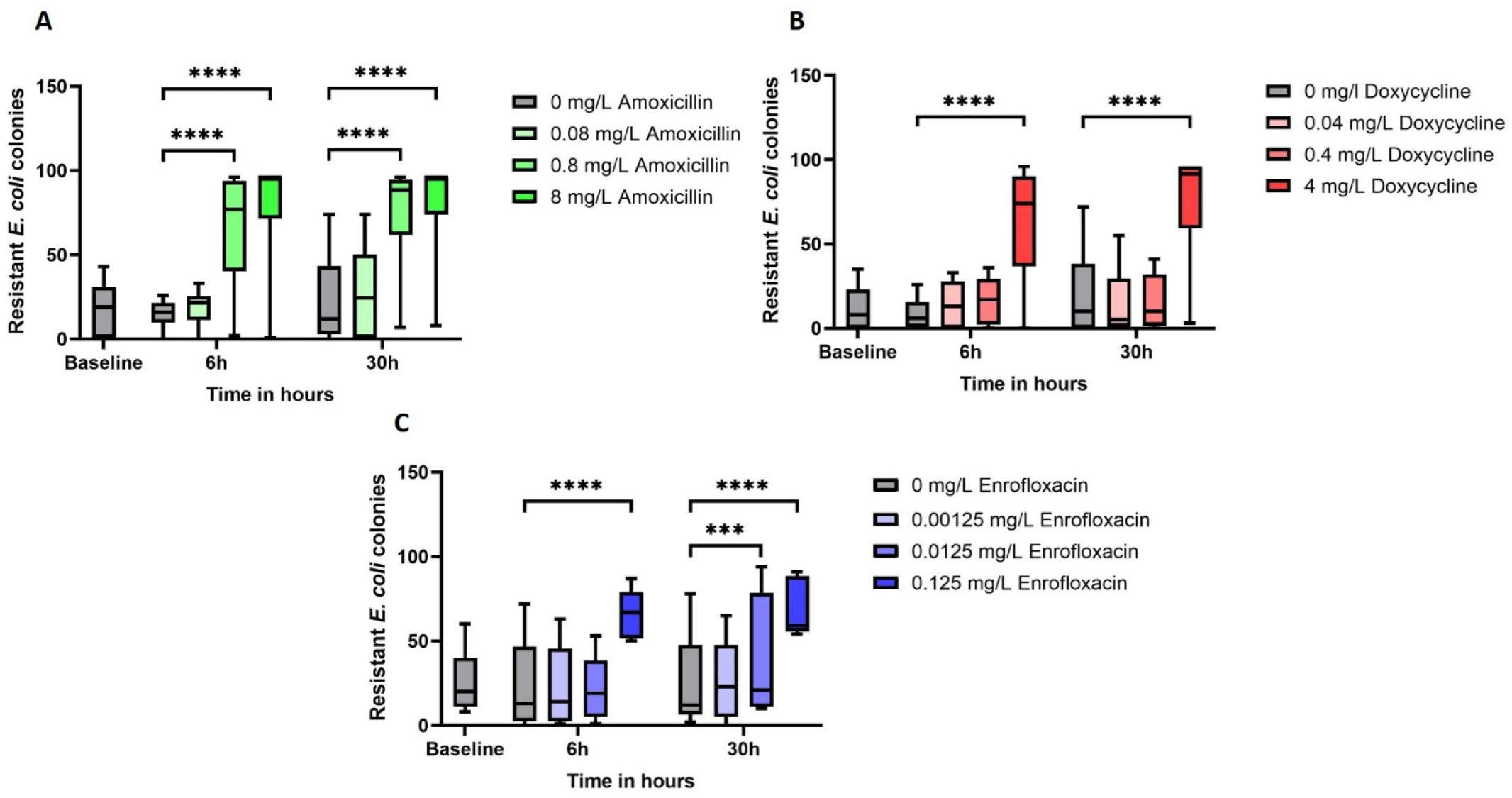
*Resistant* E. coli *isolates from the caecal fermentation. In the graphs the resistant colony count is shown for the different treatments. **A** Shows the amoxicillin treatment, **B** shows the doxycycline treatment and **C** shows the enrofloxacin treatment. In the box and whiskers the median and the minimal and maximum values are displayed. *** p ≤ 0.001, **** p ≤ 0.0001*

### Resistome analysis of the caecal fermentations

We analysed the resistome to investigate the effect of treatment with different concentrations on the number of resistance genes. We considered the specific increase in resistance genes that causes resistance to the antimicrobial which the caecal fermentation was treated with, shown in Figure 3.

**Figure 3.**
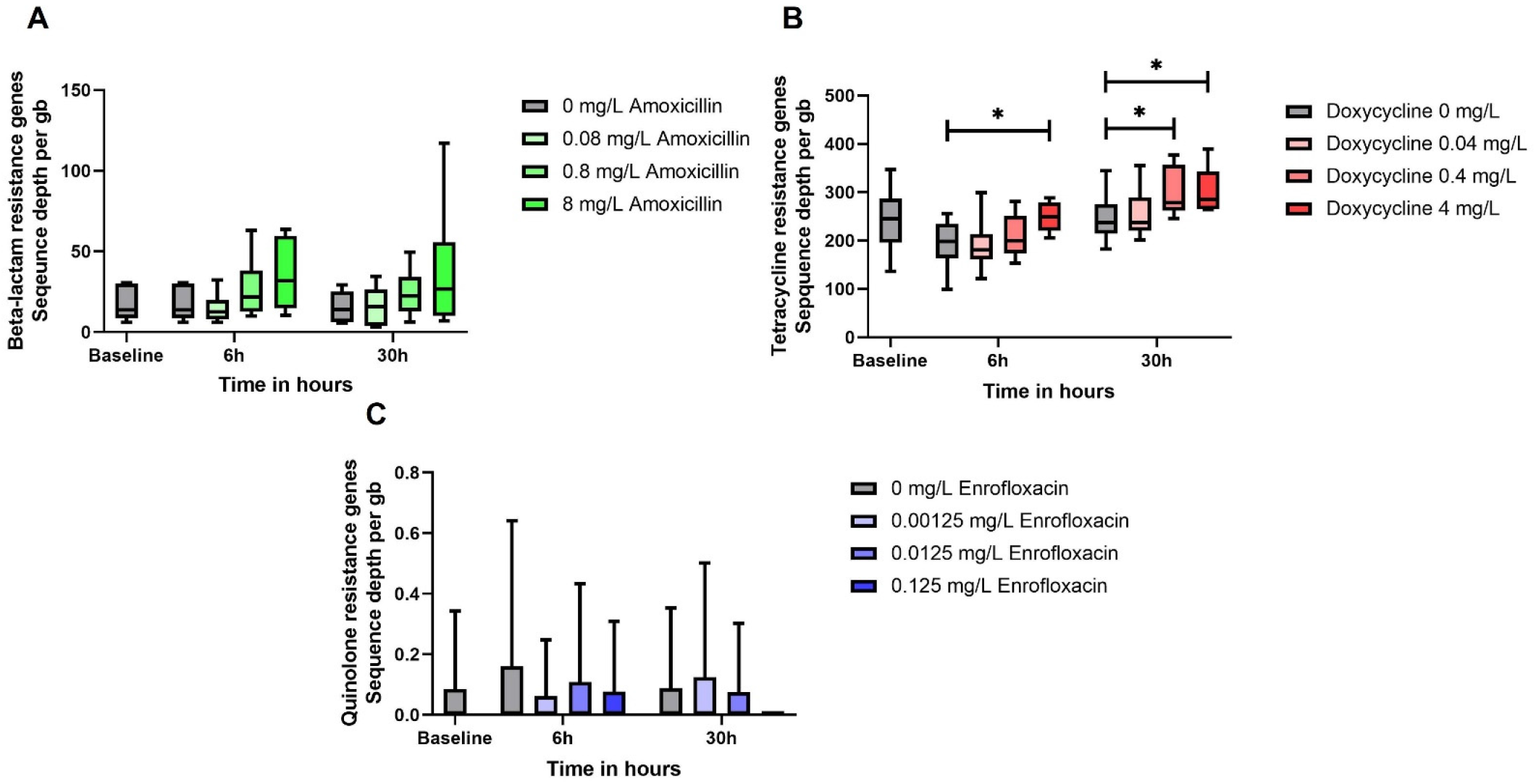
*Resistance genes per gigabase in the different treatments. **A** shows the beta-lactam genes in the samples treated with amoxicillin. The graph in figure **B** presents the tetracycline genes in the samples from the treatment with doxycycline and in graph **C** the qnr genes are shown in the treatments with enrofloxacin. In the box and whiskers the median and the minimal and maximum values are displayed. * p ≤ 0.05*

The treatment with amoxicillin did not show an increase in the number of beta-lactam genes compared to the control group. We did find more variation of the resistance depth of the beta-lactam genes per gigabase in the highest concentration of 8 mg/L however no significant difference was found. In contrast to the treatment with doxycycline where the highest concentration of 4 mg/L gave a significant difference with the control groups, bothl for the 6 hour sample (p=0.037) and for the 30 hours sample (p=0.030). In addition, the concentration of 0.4 mg/L doxycycline at timepoint 30 hours also resulted in a significant difference (p=0.050). We examined the sequence depth, number of times a nucleotide has been read, of quinolone resistance genes (*qrn*) for the treatments with enrofloxacin but no differences were found. Resistance towards enrofloxacin or fluoroquinolones is mostly mediated by SNPs in the quinolone resistance-determining regions (QRDR) in the *gyrA* and *parC* genes that play a crucial role in DNA replication (35). However, we did not determine the difference in SNPs between the control treatments and enrofloxacin.

### Microbiome composition

The microbiome was analysed to study the effect of the antimicrobial treatments on the microbial composition. We studied the effect of treatment on the alpha and beta diversity of the fermentations. Alpha diversity was determined by the Shannon index and the beta diversity was estimated by the Bray-Curtis distance from the baseline measurement (T=0) between the different concentrations within a treatment. Besides this we studied the species composition of the different treatments for a change in the microbial composition.

The different concentrations of amoxicillin did not show any differences compared to the control group in either alpha diversity or beta diversity. In addition, this was also the observation for the treatment with enrofloxacin. In contrast, the treatment with doxycycline showed a significant difference in alpha diversity in which the treatment with the concentration of 4 mg/L was different to the control at 30 hours (p=0.023). In addition to this, the beta diversity was also different between those two treatments at 30 hours (p=0.0053). Thus, the microbial composition was altered by 4 mg/L doxycycline over 30 hours where we estimated the MSC for doxycycline on the microbiome. Besides alpha and beta diversity we studied the microbial composition to observe an alteration and therefore we also analysed the relative species abundance in the caecal fermentations, shown in Figure 5. The species composition differed between the treatments and the control samples. Certain species disappeared under detection level in the microbiota and were replaced by other species. For doxycycline 4 mg/L at 6 and 30 hours we observed *Klebsiella pneumoniae* abundance which is in contrast to the control fermentation at the same timepoints. The enrofloxacin treatment resulted in less abundant *E. coli* compared to the control group especially after 30 hours. The amoxicillin treatment induced mostly a difference in the microbial composition after 6 hours since the composition after 30 hours was comparable to the 30 hours control.

**Figure 4.**
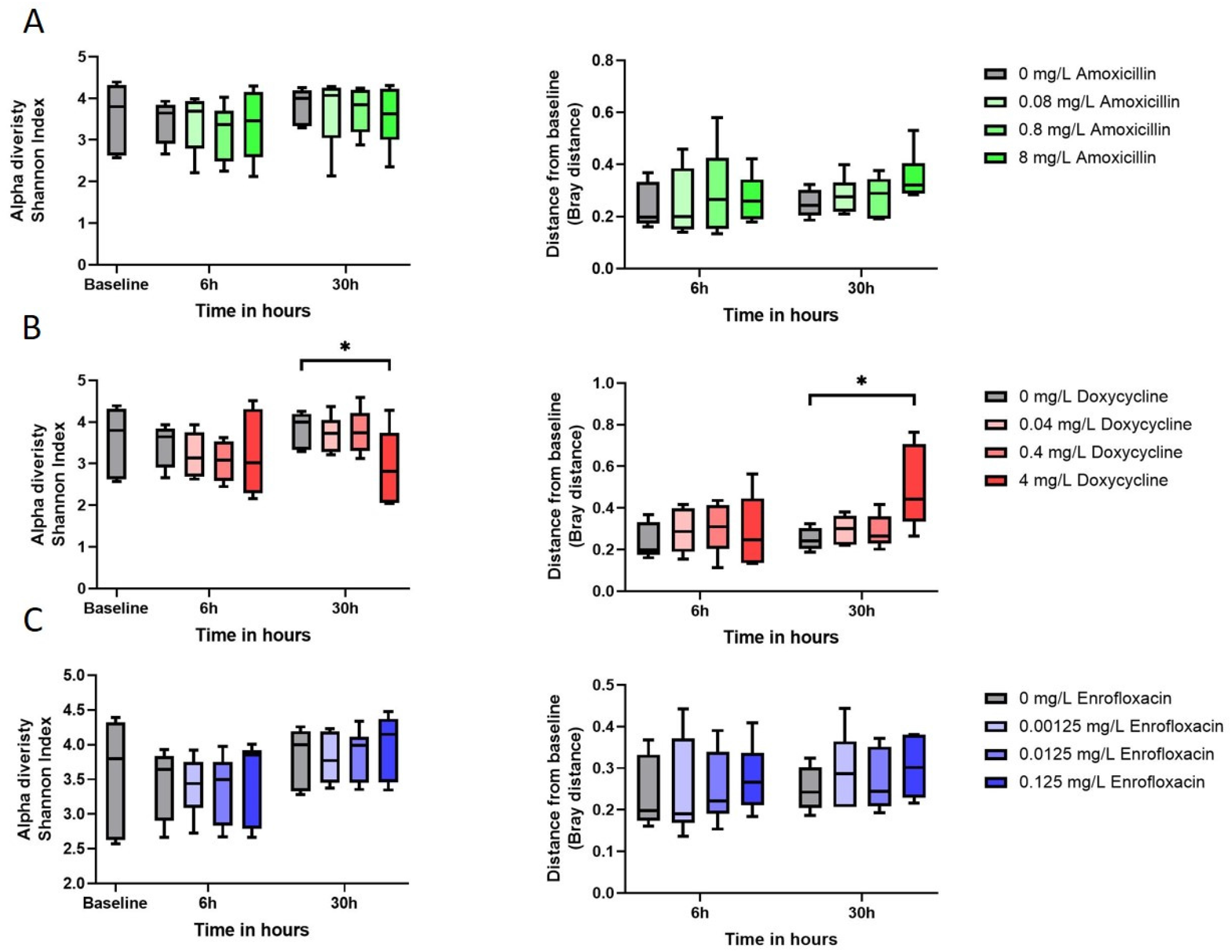
*Alpha and beta diversity of the different treatments. Alpha diversity is measured in the Shannon index and beta diversity is determined with Curtis-Bray distance. **A** the amoxicillin treatment, **B** the doxycycline treatment and **C** the treatment with enrofloxacin. In the box and whiskers the median and the minimal and maximum values are displayed. * P ≤ 0.05.*

**Figure 5.**
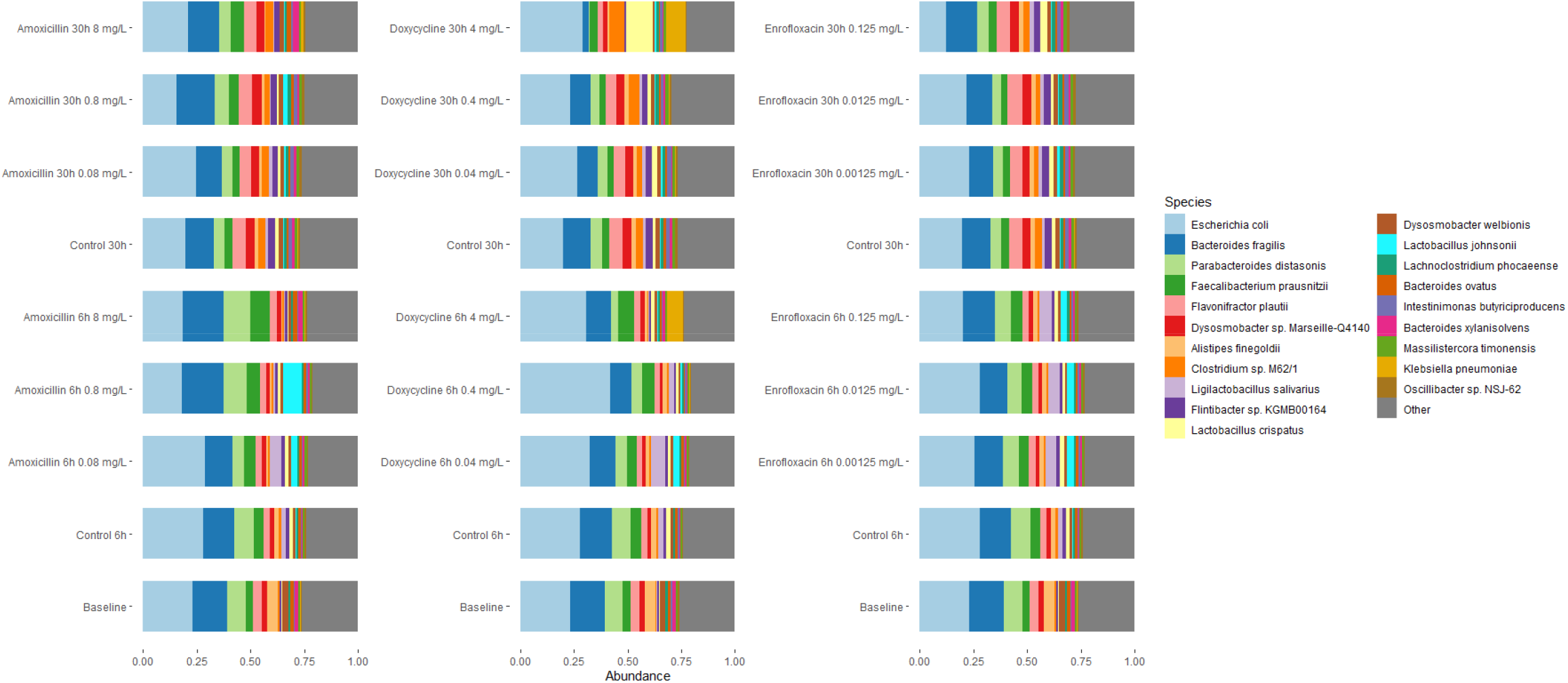
*The relative abundance in the caecal fermentations at species level. The 20 most abundant taxa are shown in the legend on the right side.*

## Discussion

The objective of this study was to determine MSCs values for amoxicillin, doxycycline and enrofloxacin in a complex microbial community by using caecal fermentation assays in which we exposed caecal contents of broilers to sub-inhibitory concentrations. Furthermore, we determined the microbiome composition in the caecal fermentations after exposure of the antimicrobial compounds. The conditions were chosen to mimic the complex physiological environment as accurately as possible. The actual MSCs are probably lower than the MSCs we determined in our experiment, since the range of concentrations we used differed by 10-fold steps. This implies that the MSC could range between the MSC determined and the concentration below that. Therefore, we display the MSCs as a range between two values. First, we conducted phenotypic resistance assays with *E. coli* which provided an estimation of the MSC from 0.08 mg/L – 0.8 mg/L for amoxicillin, 0.4 mg/L – 4 mg/L for doxycycline and 0.0125 mg/L – 0.125 mg/L for enrofloxacin. Secondly, we studied MSCs for the microbiome and resistome. Here we only found an indication of the MSC for the doxycycline treatment, since we did not find a significant difference in the amoxicillin or enrofloxacin treatment. The resistome data provided an MSC which also ranged from 0.4 mg/L to 4 mg/L and the microbiome analysis provide an estimate between 0.4 mg/L to 4 mg/L solely at 30 hours.

The most clear MSC breakpoint was observed in the analyses of phenotypic resistance of *E. coli* isolated from the caecal fermentations. We were able to distinguish an increase in the resistant *E. coli* isolates compared to the control groups per treatment. Our results gave an insight into the MSC in a complex environment for the indicator organism *E. coli* that is used to measure anthropogenic influence in the environment which underlines the value of MSC estimations (36). Moreover, in recent studies it is shown that AMR of *E. coli* is positively correlated with the AMR in the bacterial population (37, 38). The comparison of the MSC observed under defined condition in the lab versus in a complex environment is not as straightforward. Recently Klümper et al. (2019 showed that the MSC in a complex environment is higher, which is comparable to our results (39). This counterintuitive result was explained by enhanced resistance spread in a complex microbial community, as originally one could expect the MSC to be lower because of increased strain competition between species. We suspect that antimicrobial drug target alterations, such as the activity of a beta-lactamase, may also increase the MSC.

Comparing the MSC in a complex environment to the ECOFF is not easy, as although the ECOFF is determined at population level and distinguishes wildtype bacteria from non-wildtype bacteria. The actual resistance levels measured as defined by the MIC value, takes place under laboratory conditions from which the ECOFF is derived. However, the ECOFF concentration gives an indication of selective pressure of an antibiotic at population level. Therefore, we compared our MSC values we determined to the ECOFF values from EUCAST which gave an interesting perspective on selective concentrations (40). To elaborate, the MSC of amoxicillin determined for *E. coli* was between 0.08 mg/L – 0.8 mg/L which is almost 10 to 100 times lower compared to the ECOFF which has been determined at 8 mg/L. The MSCs of *E. coli* determined in our study for doxycycline (0.4 mg/L – 4 mg/L) and enrofloxacin (0.0125 mg/L – 0.125 mg/L) are 0 to 10 times lower compared to the ECOFFs (doxycycline 4 mg/L (since 27-02-2023 8 mg/L) and enrofloxacin 0.125 mg/L) of EUCAST (40). Suggesting that selective concentrations can be up to 100 times lower than the ECOFF and still select for resistant *E. coli*. Nevertheless, *E. coli* represents a small part of the whole microbial composition which makes it therefore imperative to investigate the resistance genes in the resistome in order to include the other species (33).

For that reason, the resistome was determined in the different caecal fermentations which showed contrasting results compared to the phenotypic resistance analysis. We found an increase in the tetracycline resistance genes in the doxycycline treatment at concentrations between 0.4 mg/L and 4 mg/L after 6 hours and after 30 hours it was decreased to 0.04 mg/L and 0.4 mg/L. This was 10-fold lower than the MSC determined by phenotypic resistance for *E. coli*. This can be due to the increased pool of genes when determining the resistome compared to solely *E. coli*. In the amoxicillin treatment we observed no significant differences. We did observe an increase at the 8 mg/L amoxicillin after 30 hours, however this increase was probably a consequence of extensive variation due to the repeated experiments. However, other factors could have influenced the caecal fermentations with the amoxicillin treatment as a result of the poor stability of amoxicillin (8). Amoxicillin is temperature sensitive and since we incubated the caecal fermentations at 41 °C its stability might have decrease during the fermentation (41). In addition, the existing resistance could also influence the effectiveness of amoxicillin, since beta-lactamase is able to hydrolyse the beta-lactam ring and therefore inactivate the compound (42). Similarly, we did not observe any increase in the quinolone resistance genes in the enrofloxacin treatments but this is probably caused by the resistant mutation selection mechanism in the QRDR region in the form of point mutations (35, which are not picked up by measuring sequencing depths of transferable resistance genes.

Antimicrobial compounds can also induce a change in the microbial composition due to its selective properties (43). We did not observe alterations in the alpha or beta diversity in the treatments with amoxicillin and enrofloxacin which is conflicting with a previous study where change in the diversity was mentioned when treating with amoxicillin (11 mg/kg per kg bodyweight) and enrofloxacin (5 mg/kg per kg bodyweight) (44). The concentrations used in these studies were therapeutic and comparable to prescriptions of antimicrobials and therefore much higher than the concentrations used in our MSC study (45–47). Another reason could be that selection of bacteria in the microbiome is driven by the presence of resistance genes in the pool genes which is extensively larger when treating a flock of broilers. Antimicrobial concentrations will select only for bacteria in the composition which harbours resistant genes for that specific antimicrobial compound (48). Only the treatment with doxycycline 4 mg/L at 30 hours showed significant differences which is similar to studies with doxycycline (100 mg/mL per kg bodyweight) and chlortetracycline (2 g/L) albeit these concentrations were considerably higher than the concentrations used in our study (49, 50). The significant observation at 30 hours could be attributed to the time the microbiome needs to grow by binary fission since only a proportion of the initial microbiome survives when adding the antimicrobial (43, 51).

Another essential point could be the selective properties of the VL medium which has been used to cultivate the bacteria. It is not unlikely that not all bacterial species will survive in VL medium as bacteria have unique growth conditions and thus selection for a number of species occurs which might influence the diversity (52). The caecum contains a mixture of anaerobic and facultative anaerobic bacteria (53, 54) and the transfer of the caecum material to the Eppendorf tubes for the fermentation could cause selection since some strictly anaerobes may not have survived the brief exposure to oxygen during transport from the lab to the anaerobic hood and this might have affected the diversity. Alteration, reduction or selection of the species in the microbiome after an antimicrobial treatment has been demonstrated in the literature as we also observed (48, 55).

In this research we determined the MSC based on the assay with *E. coli*, within the resistome and the microbial composition of the fermented caecal microbiota. The conditions we used in the caecal fermentations have been set to mimic the environment in the intestine as best as possible and therefore generated, to our belief, MSCs that are applicable to the physiological situation. However, despite the optimal conditions we implemented in this study it lacks some factors compared with the natural gut environment. For example, external factors are not taken into consideration such as transmission of microbes via litter uptake, fluctuating abiotic conditions as temperature or as previously mentioned selection through the medium (52, 56). Furthermore, each repetition is from two different broilers originating from different flocks. It was unknown if the broiler flocks were treated with antimicrobial compounds. This has a consequence that each repetition consists of a different microbial flora and resistant gene pool due to other environmental circumstances the broilers encountered. This is clearly visible in our data in which we observed some extensive variation at some timepoints of the treatments. For instance, in the phenotypic resistance assays of *E. coli*, resistome analysis of the amoxicillin treatment or the diversity of the enrofloxacin treatment. Interestingly, the MSC is fairly constant.

In previous studies the MSC has been determined for a complex microbial community but in a contrasting study designs that also generated MSCs for ciprofloxacin and tetracycline which are comparable to enrofloxacin and doxycycline (56, 57). In these studies, phenotypic resistance of the present bacteria and shotgun analysis for the resistome were also performed. The MSC provided by Lundström et al. (2017 and Stanton et al. (2022 of tetracycline ranges from 1 μg/mL to 10 μg/mL. By phenotypic analysis the MSC was determined at 10 μg/mL and by examination of the resistance genes of *tetA* and *tetG* at 1 μg/mL. The MSC determined here is extensively lower than the ECOFF of 8 mg/L provided by EUCAST. Comparing the MSC found for tetracycline with our MSC determined for doxycycline is there an extensive difference since the MSC we provided was 0 to 10 times smaller than the ECOFF of doxycycline while the MSC Lundström et al. (2017 and Stanton et al. (2022 provided for tetracycline was 800 times smaller than the ECOFF (40, 56, 57). On the other hand, Stanton et al. (2022 also examined the MSC for ciprofloxacin by quantifying the *intI1* integrase at different concentrations at which they found an MSC at approximately 15 μg/mL which is comparable for the MSC we determined for enrofloxacin ranging from 0.0125 mg/L – 0.125 mg/L (57). Although the authors of these papers also investigated the MSC in a complex microbial community, a significant difference is that their samples were obtained from wastewater plants. Especially the difference for the MSC of tetracycline and doxycycline could be explained by the difference in composition of the wastewater which is a collection of faeces of non-clinical and clinical sources. This could indicate that the pool of resistance genes presence is larger at the start and therefore lower concentrations might already favourably affect the selection for resistant bacteria (58, 59).

Considering the results from previous studies, the MSC of an antimicrobial compound can be different due to the environmental context as bacterial interactions and metabolites have an influence on the MSC (60). In addition to this we also found that the MSC is lower (amoxicillin) or higher (doxycycline and enrofloxacin) in the more natural caecal fermentation assay than in rich medium, as is previously described by Klümper et al. (2019 (39). Nevertheless, the MSCs which have been estimated in our research can be used by veterinarians since it gives a better understanding about the selection window for AMR bacteria in the context of treatment of poultry. Especially in combination with persistent antimicrobial compounds, the MSCs is valuable to inform policy makers how to categorize antimicrobials to reduce the selection for AMR (60, 61).

To summarize we have established the MSCs for amoxicillin, doxycycline and enrofloxacin by using caecal fermentation assays in which we aimed to mimic the natural environment in the intestine of poultry. The most explicit MSC breakpoint was observed during the phenotypic resistant assay of the commensal indicator *E. coli*. The results gave an insight on the selective concentrations of the antimicrobial compounds used in this study and could therefore contribute to decreasing the upcoming threat of AMR.

## Acknowledgement

This work was supported by the Netherlands Centre for One-Health (NCOH). We would like to thank Dr. Michael S. M. Brouwer from Wageningen Bioveterinary Research for sharing his knowledge for cultivating bacteria in the VL-medium. Furthermore, we would like to thank the suppliers of the caeca material which have been used in this study.

## Data Availability

The sequence data were deposited in the European Nucleotide Archive (ENA) under study accession number PRJEB70300.

